# High throughput screening of a new fluorescent G-quadruplex ligand having telomerase inhibitory activity in human A549 cells

**DOI:** 10.1101/2021.11.27.470216

**Authors:** Sourav Ghosh, Debapriya De, Victor Banerjee, Soumyajit Biswas, Utpal Ghosh

## Abstract

Genome-wide analysis showed that putative G-quadruplex DNA structures are prevalent in the human genome. The presence of G-quadruplex structure in the telomere and promoter region of certain oncogenes inspired people to use G-quadruplex ligand as anti-cancer agents. G-quadruplex structures, stabilized by ligand at telomere are resolved by telomerase making the cancer cells resistant to G-quadruplex ligand. So, identification of a new G-quadruplex ligand having anti-telomerase activity would be a promising strategy for cancer therapy as about 85% of human cancers are telomerase positive. A set of the drug-like compounds were screened from the ZINC database randomly and 2284 ligands were chosen following Lipinski’s rule of five that were docked with five different G-quadruplex DNA sequences in idock. We screened 43 potential G-quadruplex binders using Z-score as a normalization scoring function. The compound (ZINC ID-05220992) gave the best score (average idock = −10.17 kcal/mol, average normalized idock = −3.42). We performed G4 FID assay, CD analysis to understand its binding with three different G-quadruplex DNA sequences, and checked its anti-telomerase activity in A549 cells using TRAP assay. We observed that this compound had an intrinsic fluorescence, capability to stain live cells with a blue fluorescence, and a specific affinity to only 22AG out of three different G-quadruplex DNA sequences under study. It showed cytotoxicity, good permeability to live cells, and a significant reduction of telomerase activity in human A549 cells at a very low dose. So, this compound has strong potential to be an anti-cancer drug.

**Graphical Abstract:** 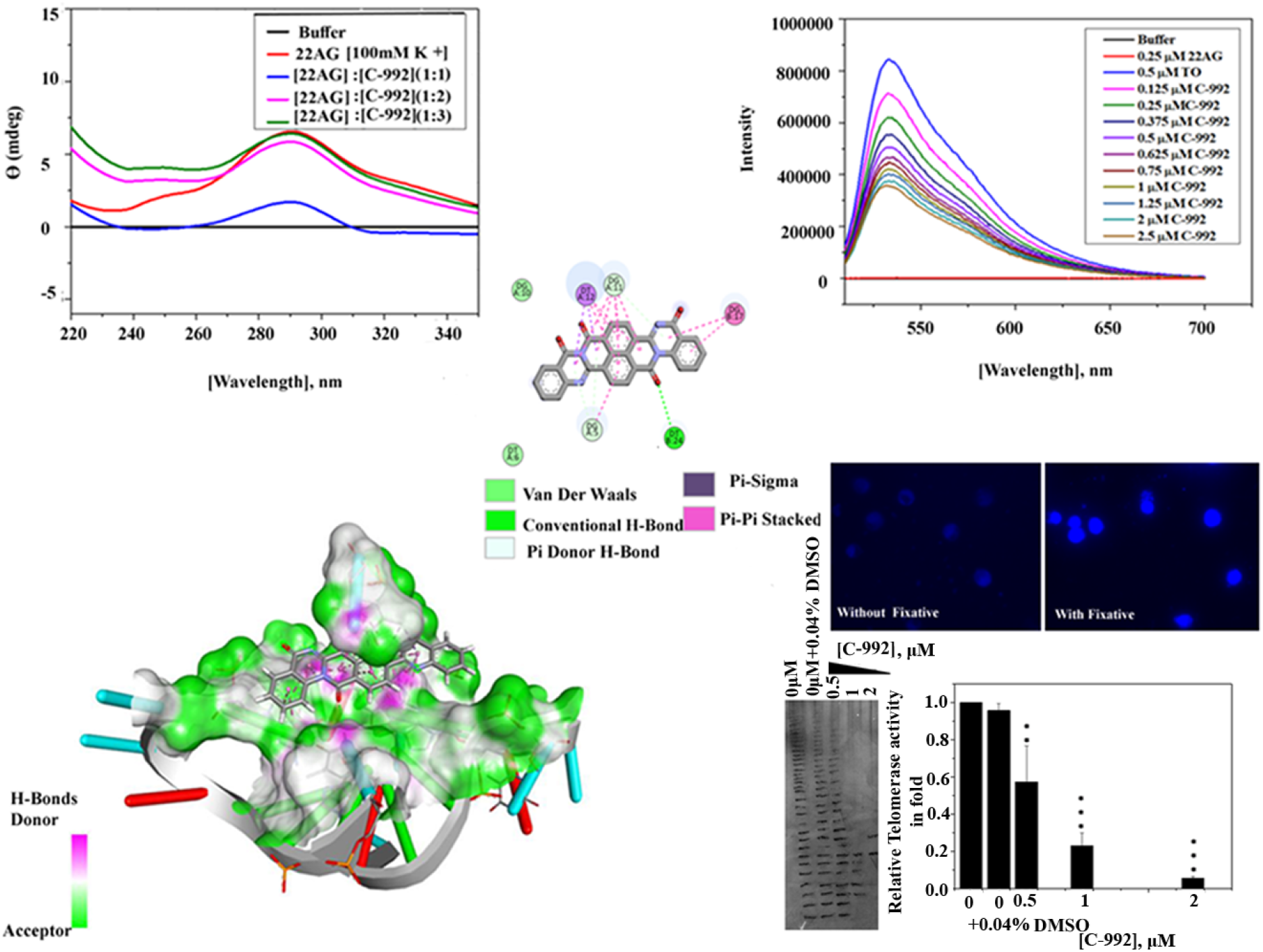

**Highlights:**

- A set of compounds were screened randomly by a High throughput method and Lipinski’s rule of five from ZINC database to identify potential G4-binders.
- The compound (ZINC ID-05220992) was screened after docking with five G-quadruplex DNA
- It binds G-quadruplex DNA 22AG as detected by TO displacement and CD spectroscopy.
- It inhibits telomerase activity in A549 cells and also cytotoxic to this cell.
- It penetrates live A549 cell in culture and stains it with blue fluorescence

## Introduction

Despite the tremendous advancement of new molecular targets like proteins and RNA for cancer therapy, DNA targets are still the mainstay of cancer treatment. Since the discovery of the G-quadruplex DNA structure, designing and searching for novel molecules targeting G-quadruplex DNA has been increased for anti-cancer therapy [1]. The G-quadruplex structure is formed when the G-rich sequence of a DNA can fold back to form a specialized three-dimensional structure where four guanines (G) are interconnected by Hoogsteen base pairing in a plane in the presence of monovalent cations like Na^+^ or K^+^ and three or more such parallel planes are stacked one after another. Either strand or both the strands of DNA of a chromosome (intra-molecular) or DNA strands from two different chromosomes (intermolecular) can form a variety of G-quadruplex structures depending on the presence of different monovalent ion concentrations, length of G-track, and length of loop interspaced G-track [2,3]. There could be a large variety of G-quadruplex DNA concerning structural and topological conformations as reviewed [4]. Detection of *in-vitro* and *in-vivo* ligand binding with various topologically polymorphic G-quadruplex DNA can effectively be monitored by measuring various fluorescence properties like quantum yield, lifetime, anisotropy of fluorescence of the ligand [5,6]. The ligand which can stabilize this G-quadruplex structure produces DNA damage, apoptosis, and cell cycle arrest when treated with cells [7,8]. The first G-quadruplex DNA structure was discovered in the telomere region [9]. Since the replenishment of telomere erosion is urgently required for highly dividing cancer cells, the interference of replenishment of telomere shortening was exploited in anti-cancer therapy using G-quadruplex ligand which inhibits telomerase to reverse the shortening of telomere and thereby induce genome instability [10–12]. Later on, it has been observed that G-quadruplex DNA is not only found in the telomere region but also distributed throughout the genome, especially at the promoter region of various oncogenes like c-MYC, c-KIT, K-RAS, hTERT, BCL-2, VEGF, HIF-1, C-MYB, PDGF-A, pRb which are responsible for cell proliferation, tumorigenesis, and anti-apoptotic effects [13]. This information makes this field more attractive for anti-cancer therapy. Even G-rich telomeric non-coding RNA (TERRA) can have various G-quadruplex structures [14]. Therefore, not only G-quadruplex DNA, G-quadruplex RNA, and G-quadruplex of DNA-RNA hybrid is also possible [4]. This makes the designing and screening of targeted G-quadruplex ligand very challenging. So far a large set of molecules have been synthesized targeting various G-quadruplex DNA structures but very few came into the clinical trial. The possible reasons are the existence of a wide variety of polymorphic G-quadruplex structures, environment-dependent DNA topology due to which *in-vitro* studies do not match with *in-vivo* studies, lack of knowledge of *in-vivo* conformation of G-quadruplex DNA, and ability of telomerase enzyme to resolve G-quadruplex DNA at telomere stabilized by ligand [15–17], etc. Therefore, identification of G-quadruplex ligand having good anti-telomerase activity *in-vivo* would be a more effective strategy for cancer therapy.

Here, we report a new G-quadruplex ligand (ZINC ID: 05220992) screened from the ZINC database based on Lipinski’s rule of five and the Z-score value of its binding with several G-quadruplex DNA using docking studies. Then the binding of this compound with G-quadruplex DNA was experimentally verified. The compound has intrinsic fluorescence property and capable of fluorescence staining of living cultured human cancer cells, has a strong binding affinity with a specific G-quadruplex DNA present in telomere, potent anti-telomerase activity.

## Materials and methods

Materials: The compound ZINC ID:05220992 henceforth named C-992 was purchased from MolProt, Latvia. Thiazole Orange (TO) was obtained from Sigma Aldrich. Tris base, DMSO, KCl were purchased from SRL India. Three different G-quadruplex DNA sequences such as 22AG (5’-AGGGTTAGGGTTAGGGTTAGGG-3’), Telo24 (5’-TAGGGTTAGGGTTAGGGTTAGGGT-3’), Telo28 (5’-TTAGGGTTTAGGGTTTAGGGTTTAGGGT-3’), and the primers for TRAP assay such as CX primer (5’-CCCTTACCCTTACCCTTACCCTAA-3’), TS primer (5’-AATCCGTCGAGCAGAGTT-3’) were procured from Eurofins, India.

Calf thymus DNA (CT-DNA) used as the control was obtained from Bangalore GeNei, India. Other fine chemicals were purchased locally.

### Methods

#### Screening of new G-quadruplex ligand from ZINC database using molecular docking

Five telomeric G-quadruplex structures were selected for virtual screening targets from Protein Data Bank (PDB) as given in Table 1 [18]. Co-crystallized ligands and water atoms were removed manually from the PDB structure of these five receptor molecules and then converted to PDBQT format from PDB using AutoDockTools for docking purposes [19]. A set of compounds were screened from 78890 compounds from the ZINC database randomly. From this set, about 2284 compounds were chosen based on Lipinski’s rule and saved in SDF format [19,20]. RdKit was used to add hydrogen and energy minimization of molecules for preprocessing of ligands using Knime workflow [21]. Then Open Babel was used to converting the SDF to PDBQT format [21]. The free and open-source docking software idock 2.2.3 was used for the docking of 2284 selected molecules at the reported ligands binding site of the five target receptor G-quadruplex DNA [22]. The details of the docking parameter of idock are given in the supplementary file S1. The binding energy and binding affinity were predicted using RF-score (pKd) and idock score (kcal/mol). All the receptors were docked with 20×20×20 grid points in the x, y, and z directions at the region of the receptor where inhibitor binds as reported in PDB. The low idock score (binding energy) and high RF-score (estimation of intermolecular binding affinity) is an indication of the possible potential of the G-quadruplex ligand. For normalization of the all receptor binding affinity, we have used Z-score (standard score) ligand q to a given receptor p and was calculated using RF-score and idock scoring function by following equations:

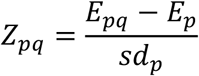

**Table-1:** Summary of iDock score and RF-Scores after binding G-quadruplex ligands with receptors G-quadruplex DNA

| PDB | iiDock score (kcal/mol) |  |  | RF-Score (pKd) |  |  |
| --- | --- | --- | --- | --- | --- | --- |
|  | Mean | Min | Max | Mean | Min | Max |
| 1NZM - 0 | -8.552371991 | -11.6 | -5.67 | 6.038148796 | 4.4 | 7.91 |
| 1NZM - 1 | -10.43320788 | -15.42 | -3.28 | 6.918490153 | 4.91 | 8.17 |
| 2HRI - 0 | -5.756652079 | -7.99 | -3.79 | 4.590310722 | 3.19 | 6.58 |
| 2HRI - 1 | -4.57854267 | -6.15 | -3.13 | 3.94854267 | 3.04 | 5.27 |
| 2JWQ - 0 | -7.402455142 | -10.61 | -5 | 5.646673961 | 3.99 | 7.34 |
| 2JWQ - 1 | -8.406617068 | -12.67 | -4.92 | 6.680866521 | 4.41 | 8.29 |
| 3CDM - 0 | -6.330568928 | -10.28 | -4.47 | 5.172984683 | 3.57 | 7.1 |
| 3CE5 - 0 | -8.565006565 | -13.3 | -6.16 | 5.70873523 | 3.93 | 7.55 |

Where E_pq_ = The idock score or RF score of ligand q docking to receptor p. E_p_ = Mean idock score or RF score value for the receptor p, sd_p_= Standard deviation of idock score or RF score of the receptor p.

A large negative value of Z_pq_ signifies the higher affinity for the ligand to the receptor. Then we categorized ligands into four groups as per Z-score values. The high G4 binder having Z-score < −2.0, the medium G4 binder having Z-score between −2.0 to −1.0, the low G4 binder having Z-score between −0.1 to 0.0, and the non-G4 binder having Z-score 0.0 to 0.1 as shown in Table 2. Finally, the compounds were sorted in ascending order according to their calculated binding energy averaged across the five receptor molecules. Discovery Studio was used for the visualization of G-quadruplex-ligands interaction.

**Table 2:**
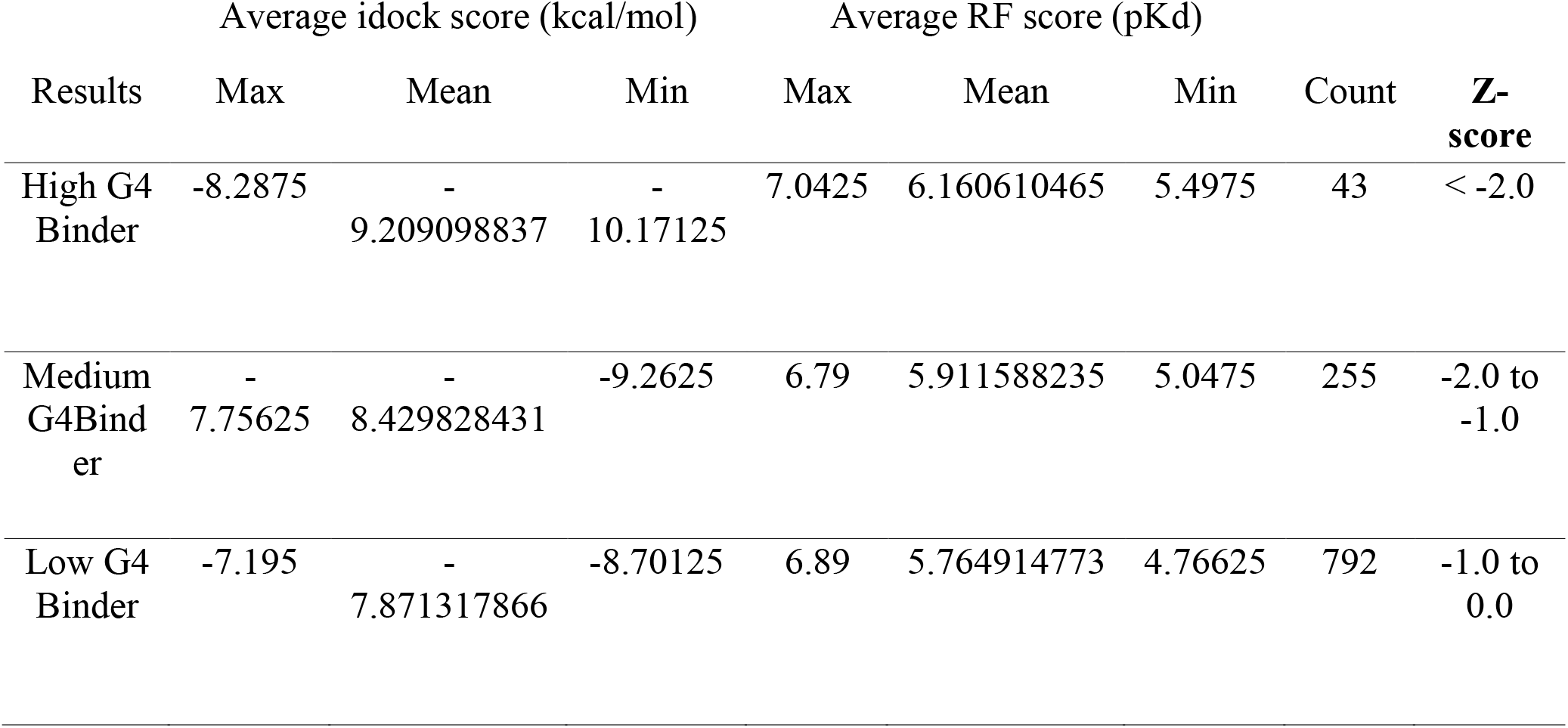
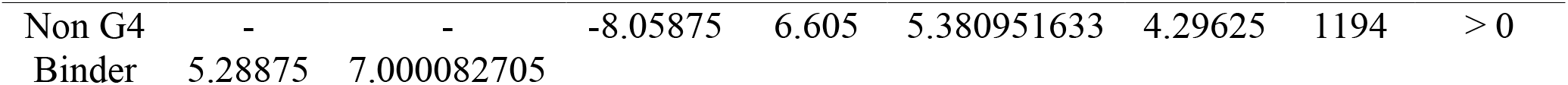
The ligands are categorized into four groups based on Z-score.

#### Cell Culture

A549 cells were obtained from NCCS, Pune and cultured in Dulbecco’s modified Eagle’s medium (HiMedia, India) supplemented with 10% heat-inactivated Fetal Bovine Serum (HiMedia, India) at 37 °C with 5% CO_2_ under humid conditions.

Preparation of solutions of G-quadruplex DNA, ligand (C-992), and Thiazole Orange (TO): The G-quadruplex DNA was dissolved in a Benchmark buffer containing 10 mM Tris-Cl, pH 7.2, and 100 mM KCl. The stock concentration was 500 μM. Then several aliquots of 50 μM in Benchmark buffer were prepared from this and stored at −20 °C for future use. Before the experiment, one aliquot was taken and heated at 95 °C for 5 min followed by chilled in ice to favor the intramolecular folding to form a specialized G-quadruplex structure. The final concentration of the DNA in the reaction mixture was 0.25 μM. A stock solution of 1 mM Thiazole Orange (TO) (Sigma, India) was prepared in DMSO. Then a sub stock of 70 μM was prepared in Benchmark buffer. The final concentration of TO in the reaction mixture was 0.5 μM for G-quadruplex DNA and 0.75 μM for CT-DNA during the G4-FID assay. Similarly, a stock solution of 5 mM of the compound C-992 was prepared in DMSO and stored at −20 °C. Then sub stocks were freshly prepared in Benchmark buffer before the experiment. The concentration range of C-992 used during titration was 0.125 μM to 2.5 μM.

### Spectroscopic method

#### G4 FID assay

The affinity of the compound C-992 towards G-quadruplex DNA can be measured by a simple fluorescence-based method known as Fluorescence Intercalator Displacement assay (FID) or G4-FID assay[23]. Here, the binding affinity of the C-992 was detected by the displacement of the fluorescent probe TO from the TO-saturated G-quadruplex DNA. The G-quadruplex DNA structure was prepared in presence of K^+^ ion as described above. TO has a strong affinity to this particular G-quadruplex structure. TO-saturated G-quadruplex DNA gives maximum fluorescence in the emission range 510 nm-750 nm with excitation at 501 nm. So, the gradual adding of C-992 to TO-saturated G-quadruplex DNA decreased fluorescence, and the percentage of TO displacement was determined by the following equation

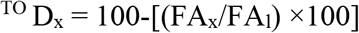

Where ^TO^Dx = Percentage of TO displacement for each concentration of compound; FA_x_ = fluorescence area of spectrum recorded after the addition of various concentrations of the ligand; FA_l_ = fluorescence area of the spectrum recorded after the addition of TO.

Intrinsic fluorescence of ligand C-992: We checked first whether the ligand C-992 has any intrinsic fluorescence property or not. First, the absorption spectra of the C-992 in Benchmark buffer were recorded in Cary Series UV-Vis spectrophotometer, (Agilent Technologies, India). Then emission spectra of the C-992 were recorded in the range 420 nm to 600 nm after excitation at its absorption maxima 370 nm in Cary Eclipse Fluorescence Spectrophotometer (Agilent Technologies, India).

Circular dichroism (CD spectra): CD spectra of G-quadruplex DNA 22AG with and without C-992 were recorded in the Applied PhotophysicChirascan spectrophotometer at a scan speed of 50 nm/min and over the range of 220 nm to 400 nm. The sample was taken in a quartz cuvette with a 2 mm path length. CD spectra of G-quadruplex in the presence of ligands (at [DNA] : [compound] ratios 1:1, 1:2 and 1:3) in Benchmark buffer (10 mM Tris-Cl, pH 7.2 and 100 mM KCl) was recorded.

Cell permeability and cell imaging by fluorescence microscope: The penetrations of C-992 in living and fixed cultured human cells were checked as per our earlier report [24]. In brief, human lung cancer A549 cells were grown on cover slips placed inside the Petri plate for 24 h. Then one set of the cover slip containing the adherent living cells was treated with 20 nM C-992 for 30 min in dark (without fixation). Another set of cover slips having adherent cells was first fixed with 3:1 methanol and glacial acetic acid at 4 °C for 2 h (with fixation). Then it was treated with the same concentration of 20 nM of C-992 for 30 min in dark. Further, cells after dislodgement from the plate by trypsinization were also tested for permeability experiment. A set of cells were dislodged from the plate by trypsinization divided into two groups. One group was treated with 20 nM of C-992 for 30 min in dark (without fixation). Another group was fixed first with 70% ethanol at 4 °C for 2 h (with fixation). Then both adherent and dislodged cells with and without fixation were observed under the 40X objective of Axio Scope A1 microscope (Carl Zeiss, Germany). Bright-field and fluorescence images were captured by Axio Scope A1 (data not shown).

#### MTT assay

Cell viability assay was done by standard method [25]. In brief, about 3000 cells were seeded in each well of a 96 well plate in triplicate. The cells were treated with different concentrations of compound C-992 after 24 h of cell seeding. Cells were allowed to incubate at 37 °C for 46 h in presence of C-992. Then the medium was replaced by fresh DMEM containing 0.2 mg/ml MTT and allowed to incubate in dark for a further 2 h. Then the medium was discarded and DMSO was added to it. The absorbance was measured in a Hitachi U-290 spectrophotometer at 595 nm.

Telomerase Repeat Amplification Protocol or TRAP Assay: Telomerase activity assay was performed as per our earlier report [26].

In brief, A549 cells were grown at 60-80% confluence. Then cells were treated with 0.5-2 μM of C-992 for 24 h. A control set was also made where cells remained untreated. Since the compound stock solution was prepared in DMSO, so another set was prepared by treating the cells with 0.04% DMSO for 24 h. After treatment, cells were trypsinized and washed twice with cold DEPC treated 1X PBS. Then cells were again washed with wash buffer followed by centrifugation at 3000 rpm for 2 min at 4 °C. Then the whole-cell extract was prepared using lysis buffer (0.5% CHAPS, 10 mM Tris-Cl, pH=7.5, 1 mM MgCl_2_, 1 mM EGTA, 5 mM β-mercaptoethanol, 10% glycerol) and was kept at 0 °C for 30 min. Next, the whole-cell extract was cleared by centrifugation at 13500 g for 15 min. Then, the supernatant was collected in a fresh tube. Protein content was measured by Lowry’s method.

Telomerase substrate (TS) [5’-AATCCGTCGAGCAGAGTT-3’] was allowed to extend by 0.9 μg of cell extract for 40 minutes at 30 °C temperature in TRAP reaction mix (20 mM Tris-HCl, 1.5 mM MgCl2, 78 mM KCl, 0.005% Tween-20, 1 mM EGTA) in a total volume of 25 μl. The telomeric repeat was amplified by PCR using CX primer [5’-(CCCTTA)3CCCTAA-3’]. The hot start Taq pol was activated at 95 °C for 5 min followed by 35 cycles of 30 s at 95 °C, 45 s at 50 °C, 1 min 30 s at 72 °C and final extension at 72 °C for 10 min. Then telomeric repeats were resolved in 10% non-denaturing PAGE and visualized after silver staining of the bands.

The intensity of each band in the control lane was added and considered as 100% telomerase activity. Accordingly, the bands of each lane were measured and normalized for the control lane. The experiment was repeated at least thrice and the statistical significance of the treated samples was evaluated by IBM SPSS Statistics version 21 software using ANOVA with Dunnett’s test. The p-values were denoted as ‘*’ (0.01 < p ≤ 0.05), ‘**’ (0.001 < p ≤ 0.01) and ‘***’ (p ≤ 0.001).

## Result & Discussion

Virtual screening of compounds having the potential to bind G-quadruplex DNA from ZINC database by molecular docking using idock:

As described in the method section, we started from 78890 compounds in the ZINC database and screened a subset of compounds following Lipinski’s rule of five and from this, a subset of about 2284 randomly chosen drug-like molecules was docked with five different G-quadruplex receptors DNA at specified region. A wide range of idock score (binding energy) and RF score (intermolecular binding affinity) was obtained for all five receptor molecules as given in Table 1.

For example, 1NZM has two docking sites and showed idock score range −11.6 kcal/mol to - 5.67 kcal/mol with a mean value at −8.55 kcal/mol and −15.42 kcal/mol to −3.28 kcal/mol with a mean value at −10.43 kcal/mol for site 1 and 2 respectively. The frequency distribution of all receptor’s idock score, RF score, normalized idock score, and normalized RF score are shown in the matrix position [1,1], [2,2], [3,3], and [4,4] respectively in Fig 1. The normalized idock score [3,3] and normalized RF score [4,4] indicated normalized value distribution calculated by Z-score. The upper and lower off-diagonal positions of the figure showed a correlation between four scoring methods. A high negative correlation value (−0.85) between idock score and RF score for all five receptors was obtained. To bring the scores of all five G-quadruplex DNA receptors on the same scale, we used a Z-score which normalized the score for all five receptors by making each of their mean to zero as shown in matrix position [3,3] and [4,4] in Fig 1.

**Figure 1:**
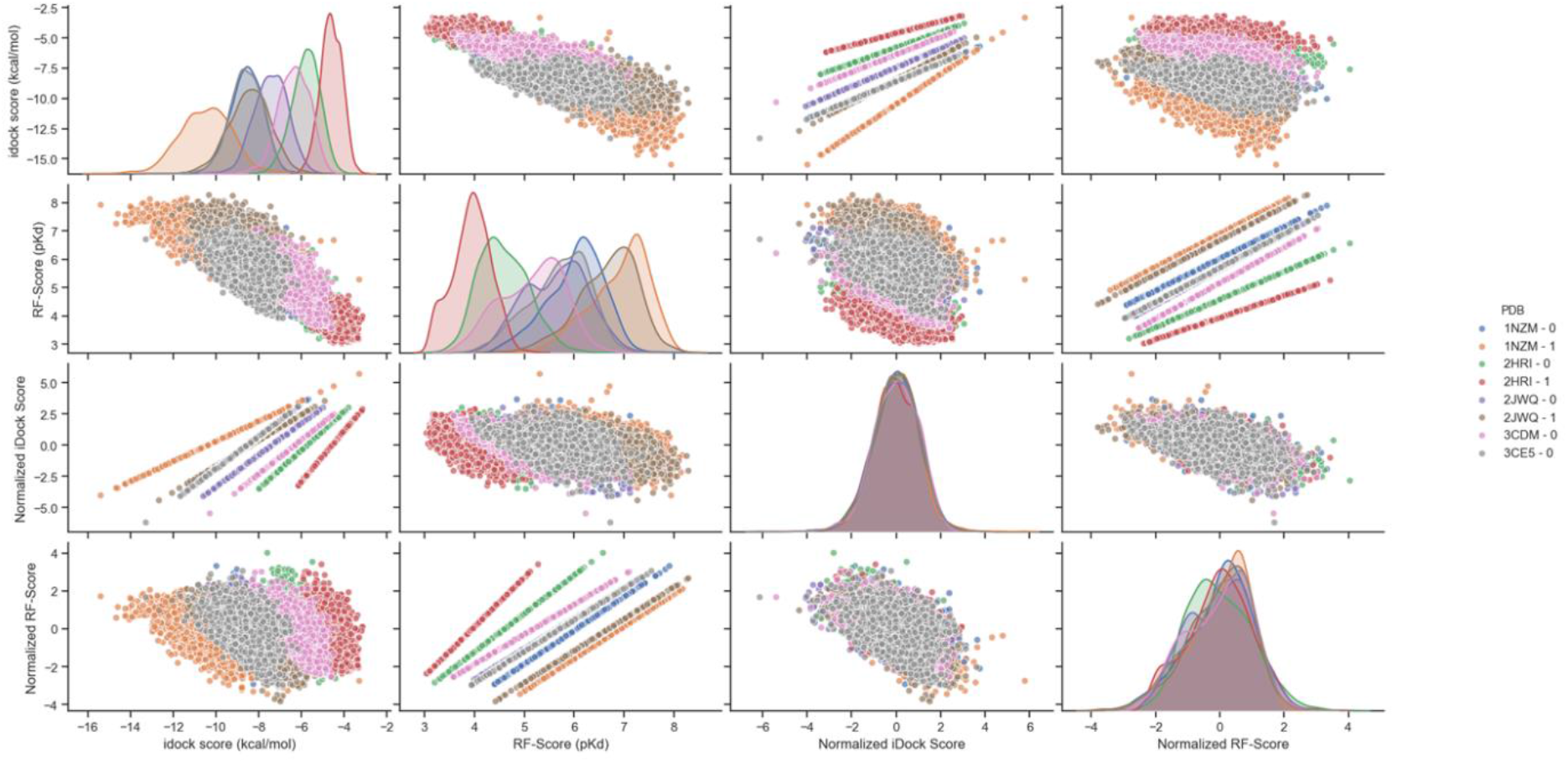
idock scores: The frequency distribution of idock score, RF score, normalized idock score, and normalized RF score of ligand-receptor interaction.

Then we grouped the ligands into four categories as described above in the method section as shown in Table 2.

About 43 ligands having Z-score less than −2.0 categorized as High G-quadruplex DNA binders (G4 binder). The frequency distribution of average idock score and average RF score for all categories is shown in matrix positions [1,1] and [2,2] respectively in Fig 2. All the 43 High G4DNA binders showed an average idock score in the range of −8.2 kcal/mol to – 10.17 kcal/mol and an average RF score in the range of 5.49 pKd to 7.04 pKd as shown in Table 2. The Scatter plot and 2D density distribution plot as shown in matrix position [2,1] and [1,2] respectively in Fig 2, distinctly indicated the distribution of four types of G-quadruplex binders with respective average idock and RF score.

**Figure 2:**
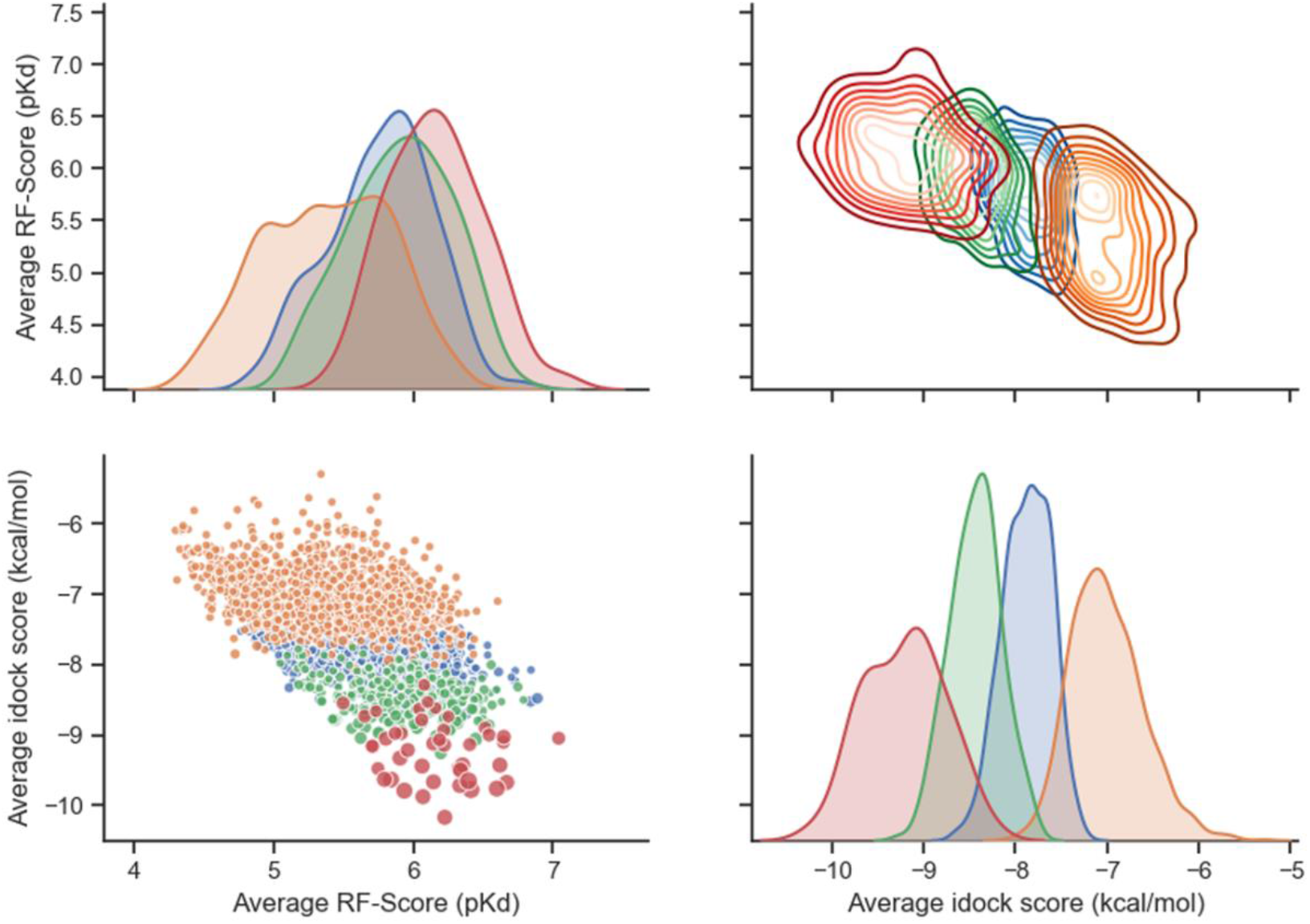
The frequency distribution of average idock score and average RF score for four category ligands. Each category ligand shows a distinct color spot or contour area.

Distinct chemical properties of the High G4DNA binder were observed among the four categories of ligands as shown graphically in Fig 3.

**Figure 3:**
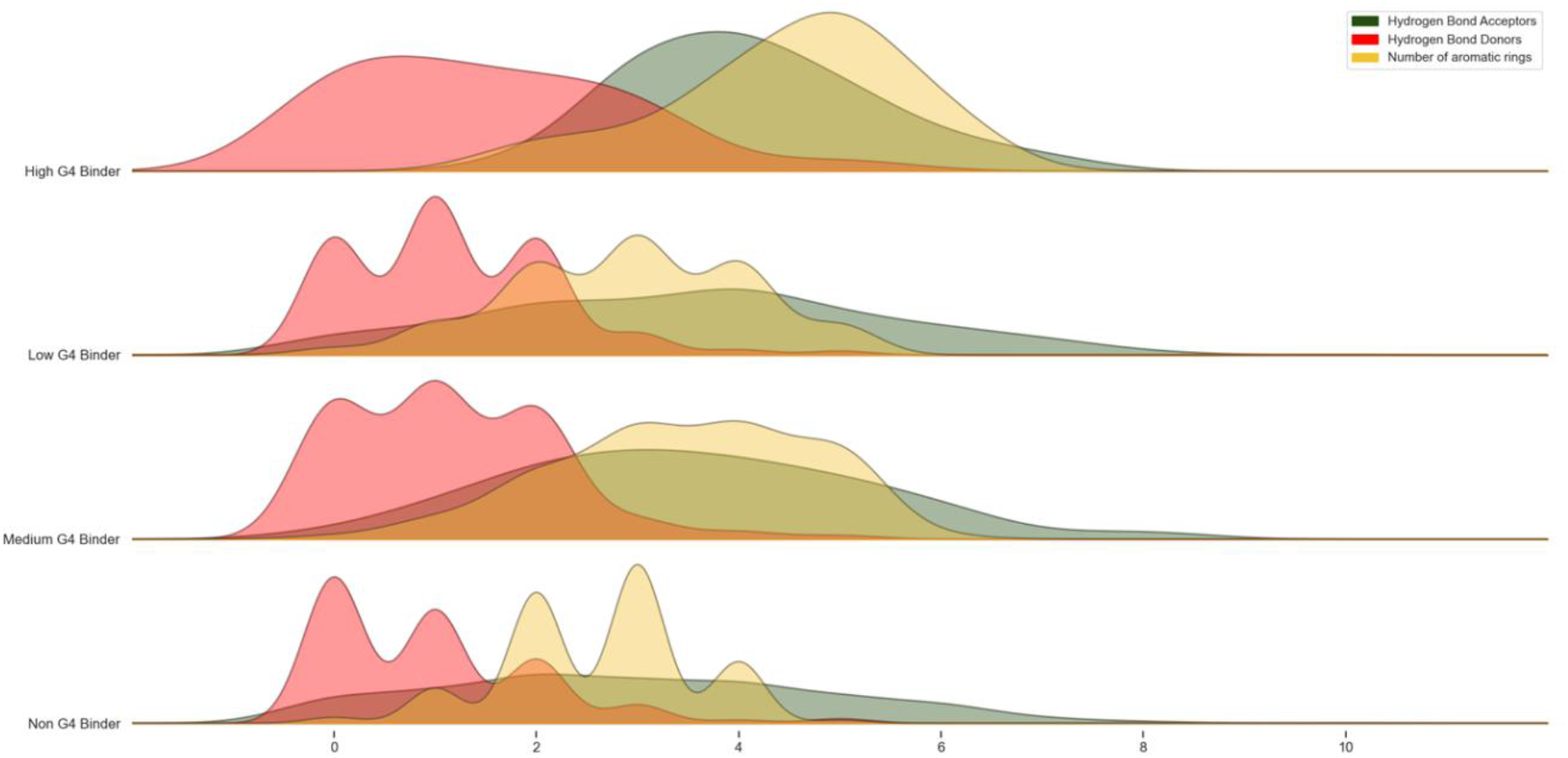
Chemical profiling of four category ligands in terms of H-bond acceptor, H-bond donor, and aromatic ring for all four category ligands.

Our result shows the presence of a high number of the aromatic ring and a comparatively higher number of H-bond acceptors than the number of H-bond donors is a prerequisite condition to be a high G4DNA binder ligand. Based on the average idock score and average RF score, three ligands such as ZINC000005220992, ZINC000013519380, and ZINC000013488698, as denoted by a circle, rhombus, and square respectively around the spot in scatter plot in matrix position [2,1] in Fig 2 were chosen. The structures of these three ligands are shown in Fig 4.

**Figure 4:**
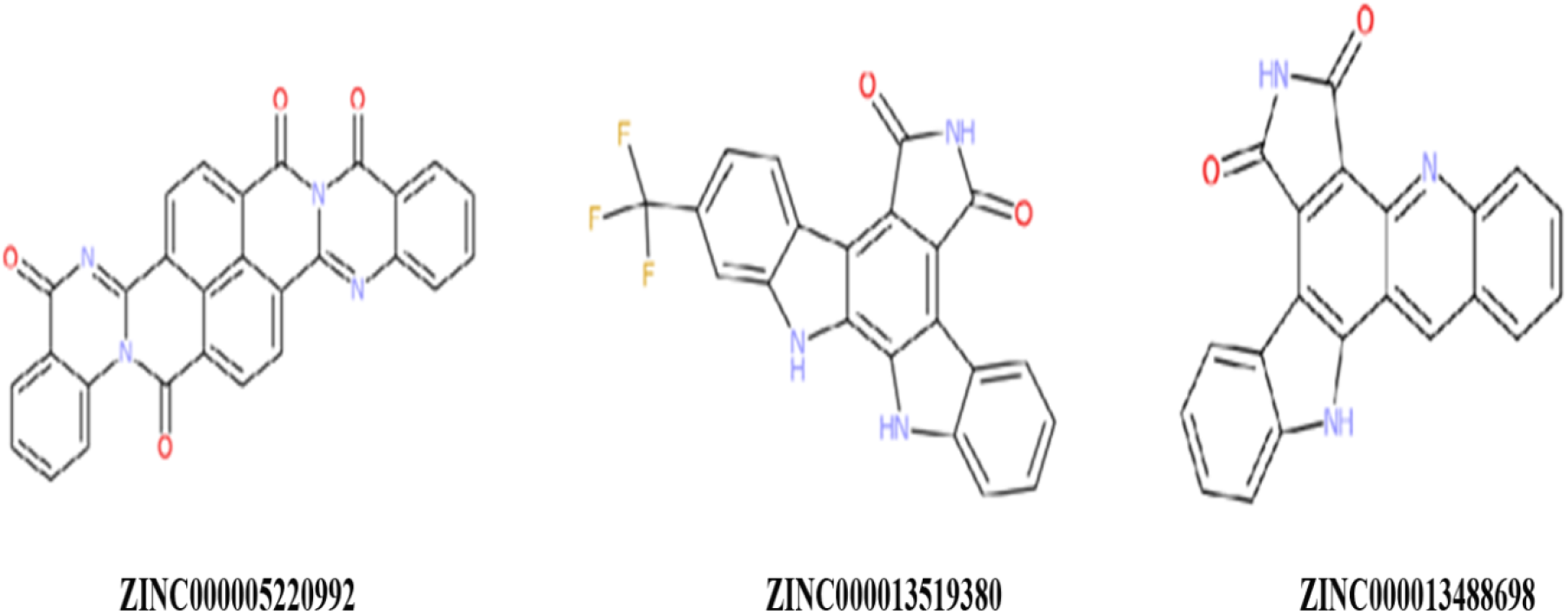
Chemical structure of three top-scored ligands. ZINC ID is given below each molecule.

The average idock score and average RF score for these ligands are (−10.17 kcal/mol, 6.22 pKd), (−9.87 kcal/mol, 6.22 pKd) and (−9.79 kcal/mol, 5.93 pKd). The best ligand as observed from the result is ZINC ID:05220992 (named as C-992) which was used for experimental validation. Several reports showed that using idock score people screened inhibitor (idock score≤ −8 kcal/mol) molecules against various receptors and worked well as per experimental validation report [27,28]. In this context, our screened compound showing a better idock score and RF score (−10.17 kcal/mol, 6.22 pKd).

The C-992 follows Lipinski’s rule of five such as molecular weight (460 Da) <500, lipophilicity measure LogP <5, hydrogen bond donor <5, hydrogen bond acceptors <10, rotatable bonds <10, and net charges = 0. The molecular docking result of C-992 with G-quadruplex DNA (PDB ID 3CE5, which is the same as 22AG except for a single base at the end) by Discovery Studio is shown in Fig 5.

**Figure 5:**
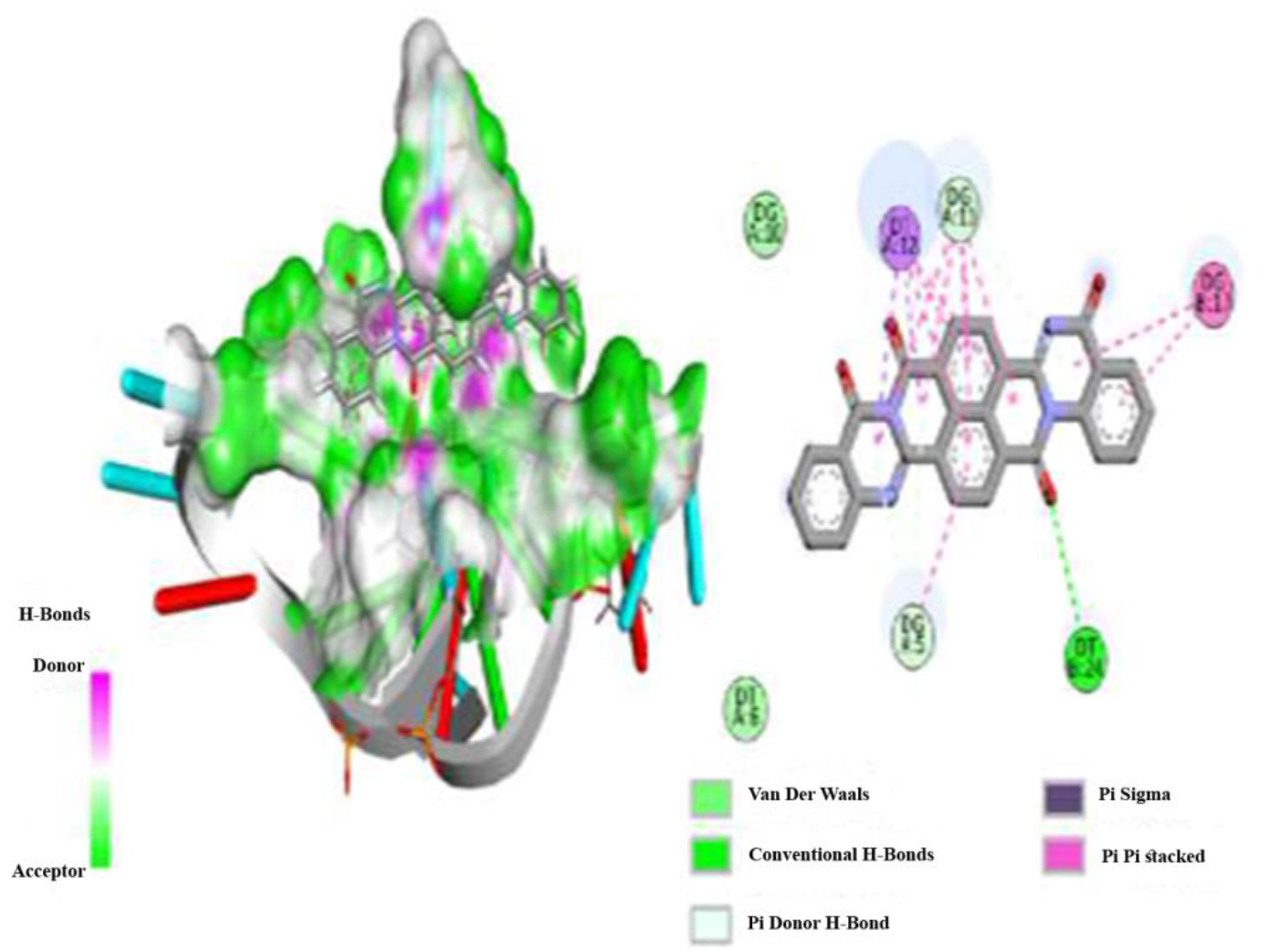
Molecular Docking results of G-quadruplex DNA (PDB ID: 3CE5) with ligand ZINC000005220992 obtained from Discovery Studio.2D interaction diagram is shown on the right side with the bond formation of ligand molecules with receptors binding sites.

The docking of this C-992 with other G-quadruplex DNA is not shown here.

### G4-FID assay

We performed the G4-FID assay to check the capability of binding of C-992 to various G-quadruplex structures. We have taken three different G-quadruplex structures like 22AG (the same as PDB 3CE5 except for single base change at the end), Telo24 (the same as PDB 3CDM except one T is absent at 5’ end), and Telo28 (PDB 1NZM, 2JWQ) as given in Table 1. Each type of G-quadruplex DNA (0.25 μM) was saturated with TO (0.5 μM) and titrated with various concentrations of C-992 to check TO replacement. We observed that 50% TO replacement (DC50) from 22AG G-quadruplex DNA occurred at 0.99 μM of C-992, whereas that from CT-DNA was occurred at 1.56 μM as evident from Fig 6a. We also observed that the DC_50_ value for Telo24 and Telo28 was more than 27.12 and 21.94 μM respectively as shown in Fig 6b.

**Figure 6:**
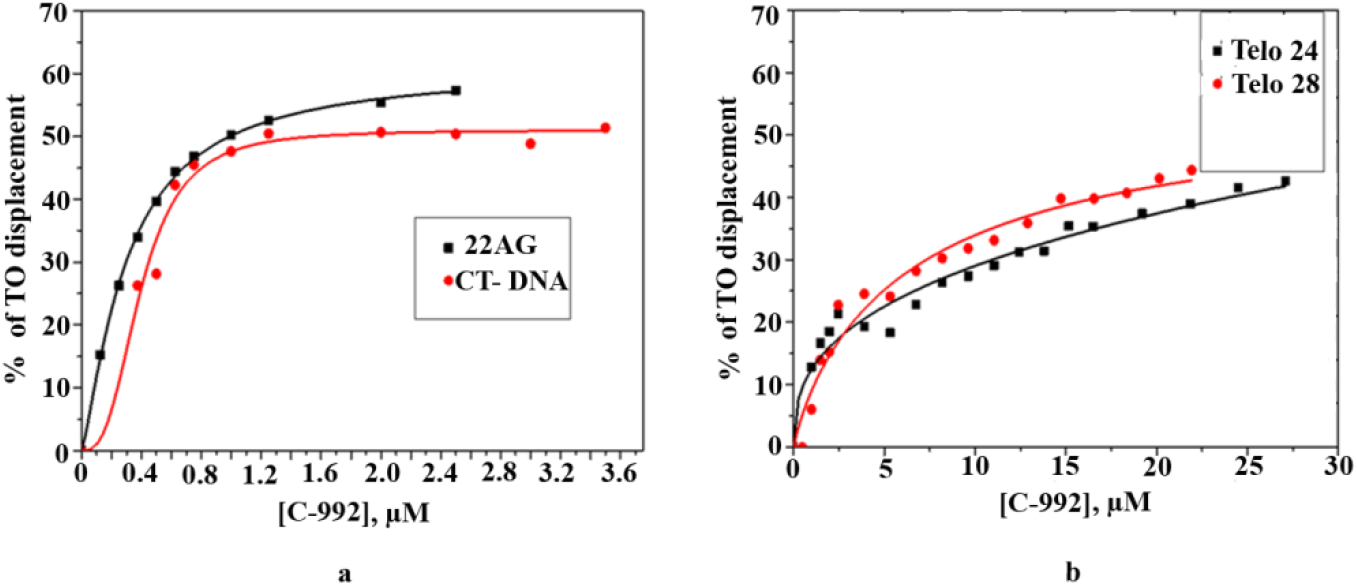
G4 FID assay: (a) TO displacement by C-992 from 22AG G-quadruplex DNA and CT-DNA; (b) TO displacement by C-992 from Telo24 and Telo28 G-quadruplex DNA.

The raw data for all these titrations are given in supplementary file S2. This data implicates that the compound C-992 has a stronger affinity to a specific G-quadruplex DNA 22AG among the three different types of G-quadruplex DNA under study. The affinity of this compound to 22AG is higher than double-stranded (CT-DNA). The 50% TO replacement (DC50 value) from various DNA by the compound C-992 is given in Table 3.

**Table 3:**
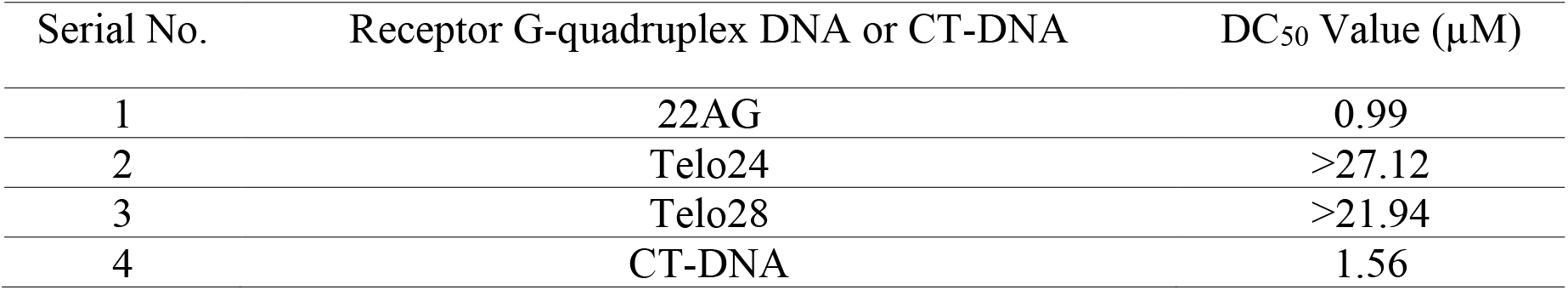
DC_50_ values of TO replacement from various receptors DNA molecules

| Serial No. | Receptor G-quadruplex DNA or CT-DNA | DC<sub>50</sub> Value (μM) |
| --- | --- | --- |
| 1 | 22AG | 0.99 |
| 2 | Telo24 | >27.12 |
| 3 | Telo28 | >21.94 |
| 4 | CT-DNA | 1.56 |

All three G-quadruplex DNA sequences have human telomeric repeats TTAGGG but they are different in the loop sequence of G-quadruplex structures. Our data implicate that loop sequence can alter binding affinity because C-992 has a very strong affinity to 22AG only out of the three G-quadruplex DNA. This observation corroborates an earlier report [29]. Moreover, the calculated DC_50_ value of C-992 for 22AG is close to reported DC_50_ values of some known G-quadruplex ligands [30,31] and even better than some of the reported compounds [32,33].

### Absorption and fluorescence emission spectra of C-992

The compound C-992 has intrinsic fluorescence property. We have taken various concentrations of C-992 (4.13-32.62 μM) in Benchmark buffer (10 mM Tris-Cl, pH 7.2 and 100 mM KCl) and scanned wavelength range from 300 nm to 600 nm to get absorption spectra as shown in Fig 7a. The maximum absorption was observed at 363 nm. We have excited our compound at a different wavelength (320 nm - 370 nm) and fluorescence emission was recorded as shown in Fig 7b. This data suggests that the molecule has an added property – it has good intrinsic fluorescence and higher intensity was observed when excited at 320 nm.

**Figure 7:**
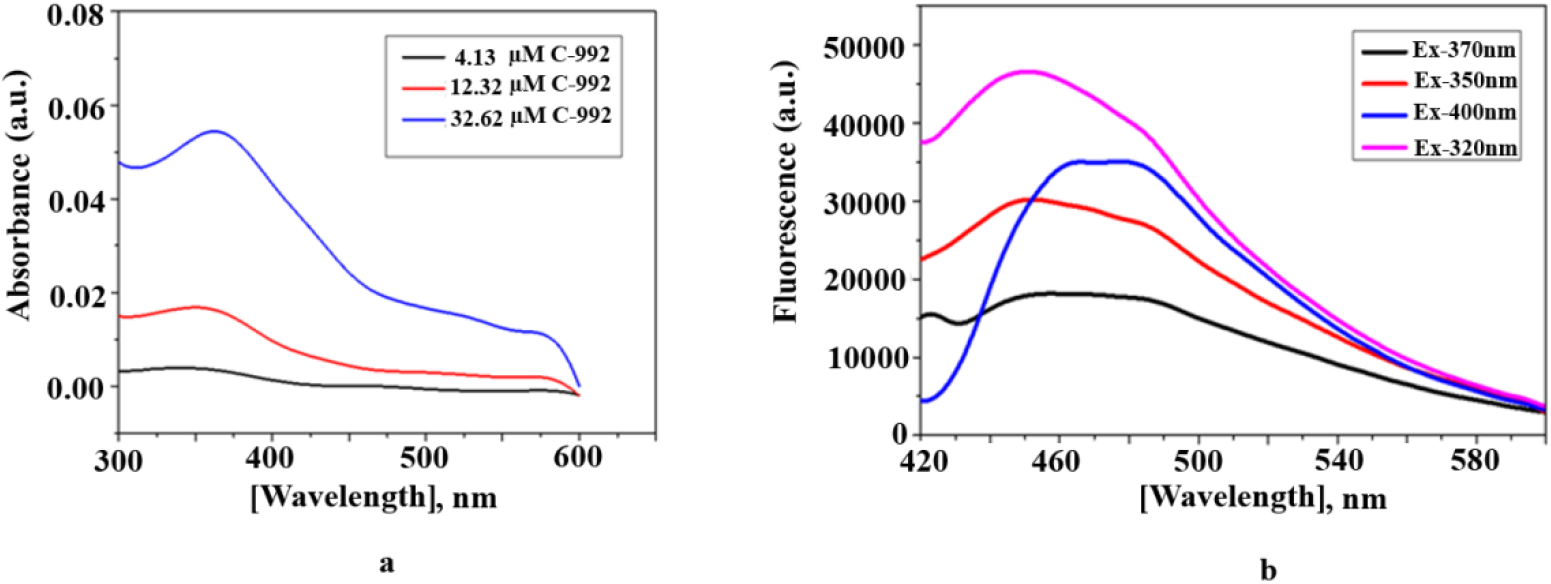
(a) Absorption spectra of different concentrations of C-992 (4.13-32.62 μM) in Benchmark buffer (10 mM Tris-Cl, pH 7.2 and 100 mM KCl); (b) intrinsic fluorescence spectra of C-992 in the same buffer with different excitation wavelength.

### CD Experiments

CD Experiment was done to determine the G-quadruplex (22AG) binding property of the compound in K^+^ containing buffer as mentioned in materials and methods. 22AG DNA forms a hybrid structure in 100 mM K^+^ containing buffer with a strong positive peak at 295 nm with a shoulder at 265 nm, a small positive peak at 250 nm, and a small negative peak at 235 nm as per earlier report [34]. Generally, 22AG takes parallel or hybrid anti-parallel structure in K^+^ buffer. In our case, it takes the same conformation as evident from the typical positive peak at 295 and 250 nm as shown in Fig 8.

**Figure 8:**
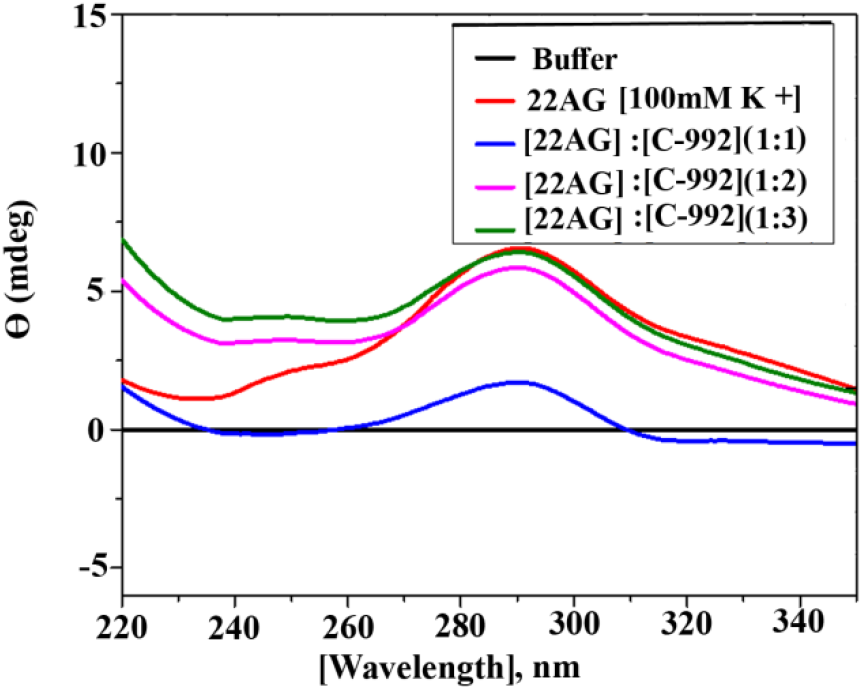
CD spectra of free 22AG and C-992 bound 22AG in Benchmark buffer (10 mM Tris-Cl, pH 7.2 and 100 mM KCl)

There were a drastic decrease in both 295 and 250 nm peaks during titration of 22AG with the C-992 at the ratio [DNA]:[compound] = 1:1. However, both the peaks dramatically increased along with an overall change in ellipticity at the ratio [DNA]:[compound] equal to 1:2 and 1:3. This data implicates that our compound C-992 has the capability of binding the G-quadruplex structure of 22AG and thereby can alter the three-dimensional conformation of the DNA.

### Cellular uptake & cell imaging by fluorescence

To observe the cellular uptake of this compound by the live-cell A549 under culture condition with and without using a fixative, we incubated the A549 cells in presence of a very low concentration (20 nM) of C-992 in PBS for 30 min at 37 °C as per our previous report [24,35,36]. Then the cell images were captured under bright field mode (data not shown) and fluorescence mode using a filter set 49 of an Axioscope A1 fluorescence microscope (excitation 365 nm and Emission 445/50). It is evident from Fig 9a that C-992 is capable of entering into cells through the live cell membrane and label the cell with sky blue fluorescence color. We also incubated the trypsinized cells in presence of C-992 after fixation with 70% ethanol. This fixation makes the cell porous and hence increases the permeability of the membrane. In addition to binding G-quadruplex DNA, it also binds double-stranded DNA with comparatively lower affinity. So, we got a brighter fluorescence image for porous cells as shown in Fig 9b.

**Figure 9:**
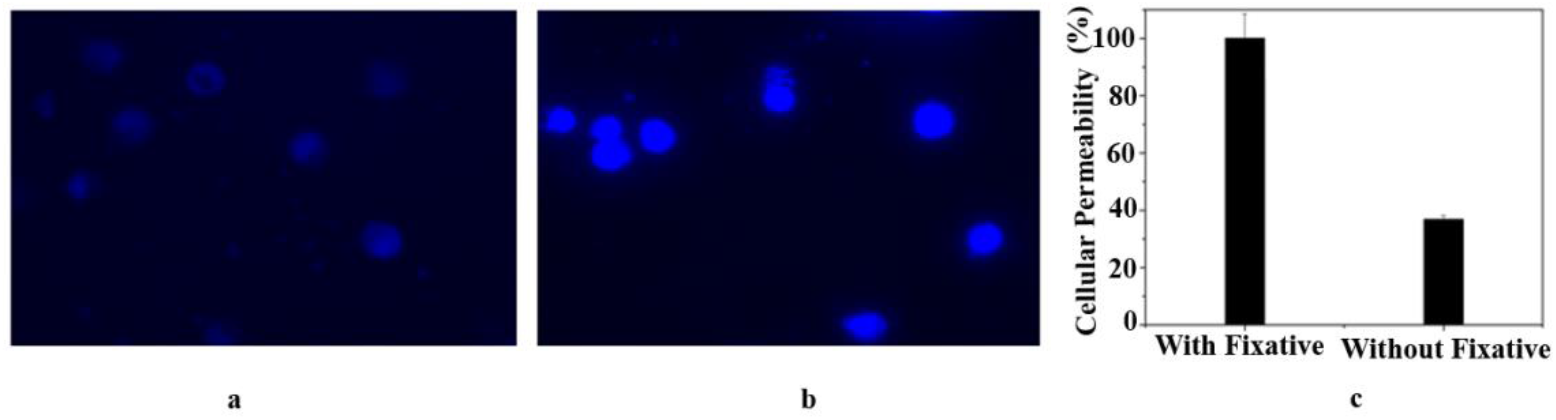
Cell permeability and live-cell imaging using C-992; A549 cell imaging under Axioscope A1 in fluorescence mode without fixation (a) and after fixation (b) of trypsinized cells; (c) percentage of cellular permeability of the compound C-992 (with and without fixative).

Taking the image intensity of the porous cells (after fixation) as 100% permeable, we calculated the intensity of cell image without fixation and observed about 40% permeability of the compound in live cells as shown in Fig 9c. We also did the permeability of the compound in A549 cells in adherent conditions and got a similar result (data not shown). Our data suggest that this compound has a strong potential to enter into a living cell and capable of lighting up the nucleus.

### Cell viability assay with MTT assay

We have checked cell viability by MTT assay. The A549 cells were treated with various doses of C-992 (10-100 μM) and incubated for 46 h. The cell cytotoxicity is low with the dose range we have used. About 87% and 65% cell viability was observed at 10 μM and 100 μM of C-992 respectively. The data is shown in Fig 10.

**Figure 10:**
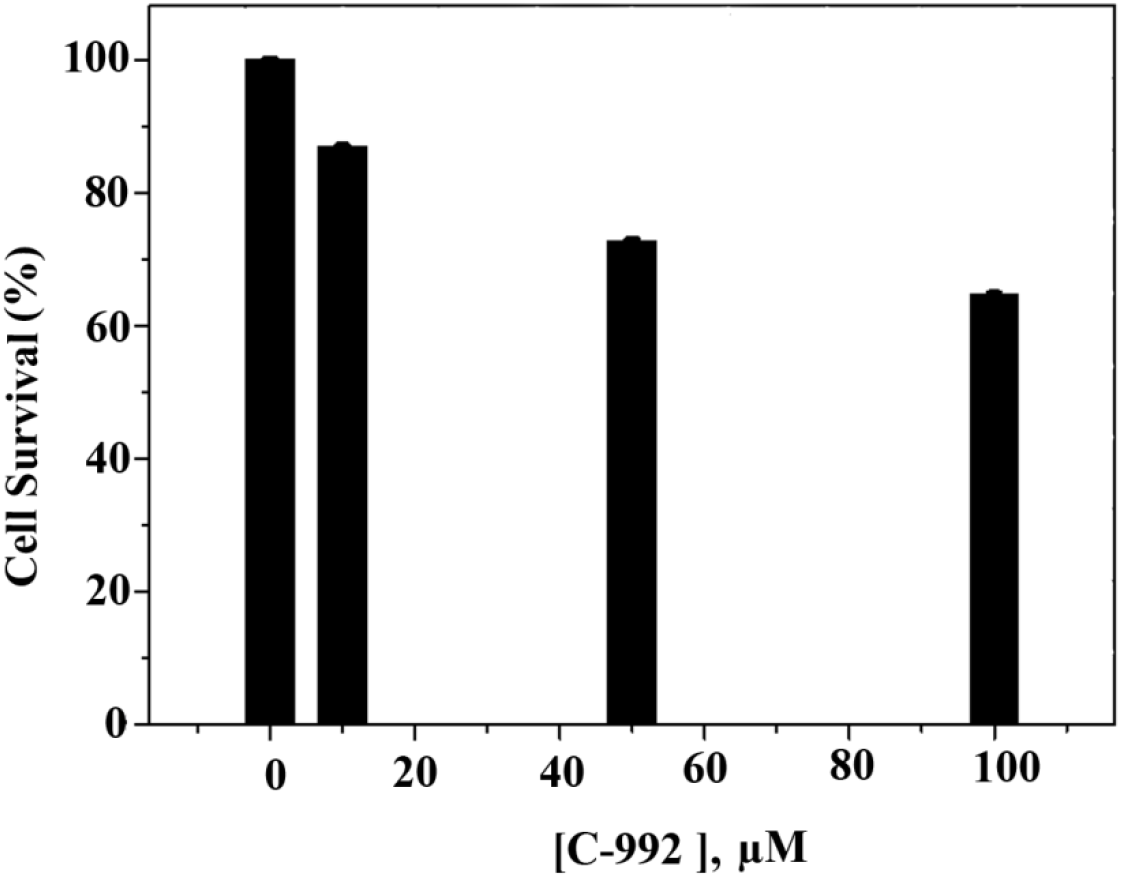
Percentage of A549 cell survival after treatment with various concentrations of C-992 for 46 h by MTT assay.

### Telomerase inhibition by C-992 in A549 cells

We observed that C-992 is a potential G-quadruplex sequence binder and it has an affinity for the telomeric G-quadruplex sequence. Hence, we measured the telomerase activity by TRAP assay. A549 cells were treated with different doses of C-992. An untreated control set and another set treated with 0.04% DMSO were also taken into consideration because the highest dose of the compound contains 0.04% DMSO in culture media. There was no difference in the number of bands between the untreated control and 0.04% DMSO sets (Fig 11a). This proves that 0.04% DMSO does not affect telomerase activity. However, a dose-dependent reduction of telomerase activity was observed in A549 cells treated with C-992 (0 – 2 μM) as shown in Fig 11a.

**Figure 11:**
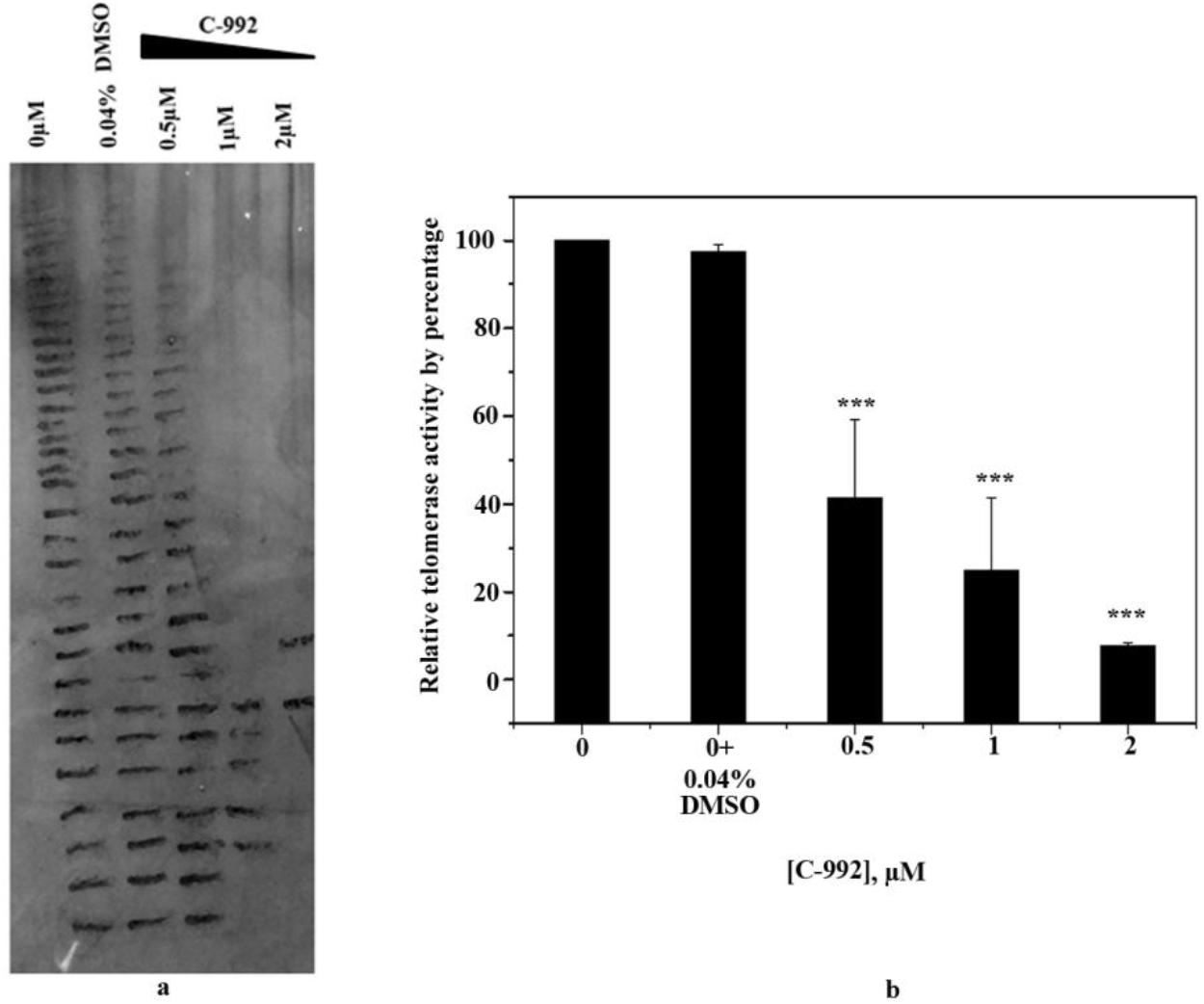
TRAP assay of A549 with C-992. a.Gel image shows C-992 can inhibit telomerase activity in a dose-dependent manner in A549 cells. b. Quantification of TRAP assay in the 11a. Student T-test was performed, p values are as follows ‘*’ (0.01 < p ≤ 0.05), ‘**’ (0.001 < p ≤ 0.01) and ‘***’ (p ≤ 0.001).

Notably, a significant (0.001 < p ≤ 0.01) reduction of telomerase activity was observed at a dose as low as 0.5 μM as shown in Fig 11b. The telomerase activity further diminished below 10% at a dose of 2 μM. This data implicates that C-992 is a potent telomerase inhibitor. Several reports showed that the G-quadruplex ligand can inhibit telomerase activity [35–40]. The proposed mechanism by which the G-quadruplex ligand inhibits telomerase is by stabilizing the G-quadruplex sequence of the telomere, by stabilizing the G-quadruplex sequence of the promoter region of hTERT or reduction of hTERT expression and by direct interaction with the catalytic domain of hTERT [41,42].

A different strategy was opted to investigate the actual mechanism of telomerase inhibition; moreover, it was required to determine whether inhibition of Taq DNA pol is responsible for the TRAP assay result. So, the same TRAP assay was performed with control A549 cell lysate where the cells were not treated with C-992. C-992 was added at a concentration of 2 μM to the TRAP reaction mix during the reaction since it was the highest dose of C-992 used for the earlier TRAP assay (Fig 11). A control set was also prepared where C-992 was not added and another set was prepared with 0.2% DMSO since C-992 was dissolved in DMSO. After the polyacrylamide gel electrophoresis, the same number of bands were observed in each lane. Moreover, there was not much difference in the total intensity of the bands (Fig 12).

**Fig 12:**
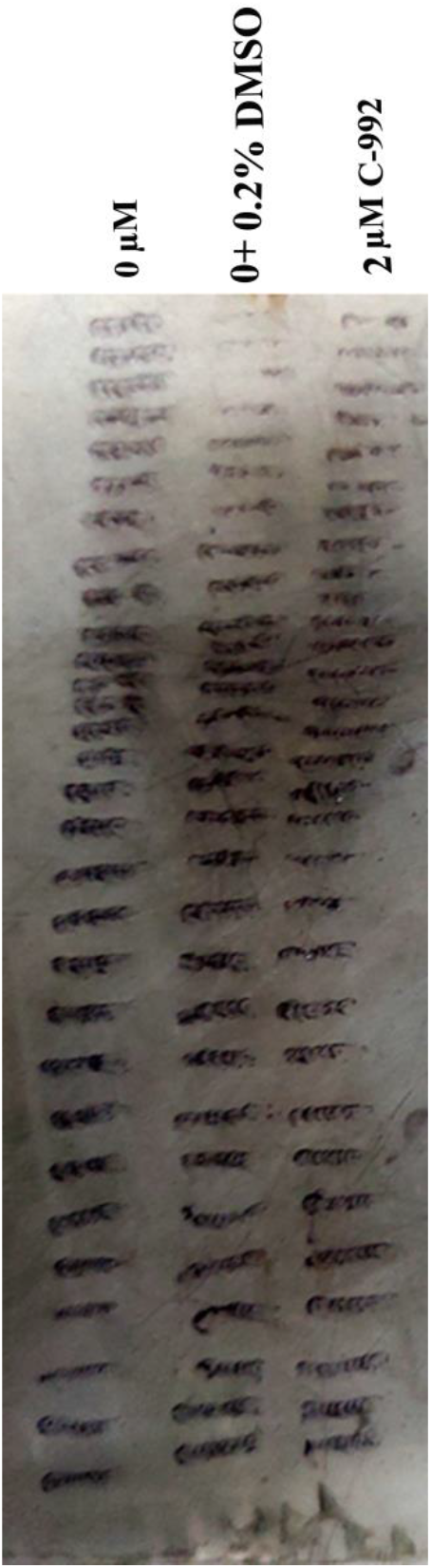
TRAP assay of A549 was performed with indicated concentration where C-992 was added to the cell-free system. The result showed no inhibition of telomerase activity.

We hypothesized that if no band was observed in C-992 lane it might be due to the formation of the G-quadruplex structure in the cell-free system or due to the inhibition of Taq DNA polymerase by the compound C-992. On the contrary to our hypothesis, bands were observed in each lane. This result is contradictory to our earlier result where telomerase activity was almost inhibited by 2 μM C-992 (Fig 11). The possible reason behind this contradictory result is that 2 μM C-992 was not sufficient to form or stabilize a G-quadruplex structure in the cell-free system and also could not inhibit Taq DNA pol activity. Whatever be the mechanism of telomerase inhibition by C-992, it is evident from our whole data that C-992 is a very good telomerase inhibitor and it can not affect the activity of Taq DNA pol.

### Conclusion

In conclusion, we report that compound C-992 from the ZINC database has a strong binding affinity with a specific G-quadruplex structure present in the telomere. Compound C-992 complies with Lipinski’s rule of five and is permeable to live cells implicating its potential to be a drug. It inhibits telomerase activity significantly *in-vivo* at a very low dose and hence it could be promising in cancer therapy. It has an added intrinsic fluorescence property and can stain the cell with blue color fluorescence.

## Funding Sources

This work was supported by UGC-SAP (DRS-II). UG thank ICMR (ICMR-SRF No.3/2/2/36/2018/Online OncoFship/NCD-III and No.3/1/3/JRF-2018/HRD-055(66816)) and DST (DST/INSPIRE Fellowship/2014/37), for providing the manpower for this work.

## Supporting Information

**S1:** Details of docking parameter of receptor G-quadruplex DNA

S1: Details of docking parameter of receptor G-quadruplex DNA
| PDB | center_x | center_y | center_z | size_x | size_y | size_z |
| --- | --- | --- | --- | --- | --- | --- |
| receptor |  |  |  |  |  |  |
| 1NZM_0 | 29.171186 | 35.135953 | 18.012209 | 20 | 20 | 20 |
| 1NZM_1 | 27.728512 | 29.371535 | 30.69614 | 20 | 20 | 20 |
| 2HRI_0 | -18.602135 | -11.650558 | -15.250596 | 20 | 20 | 20 |
| 2HRI_1 | -9.872596 | 0.1565 | -0.082 | 20 | 20 | 20 |
| 2JWQ_0 | -5.531532 | -2.952129 | 21.318581 | 20 | 20 | 20 |
| 2JWQ_1 | -4.194823 | 1.860935 | 9.099758 | 20 | 20 | 20 |
| 3CDM_0 | 2.629881 | 0.820333 | -14.777619 | 20 | 20 | 20 |
| 3CE5_0 | 17.984886 | 18.087727 | 9.447205 | 20 | 20 | 20 |

S2: Raw spectrum of TO displacement from three G-quadruplex DNA and CT-DNA.

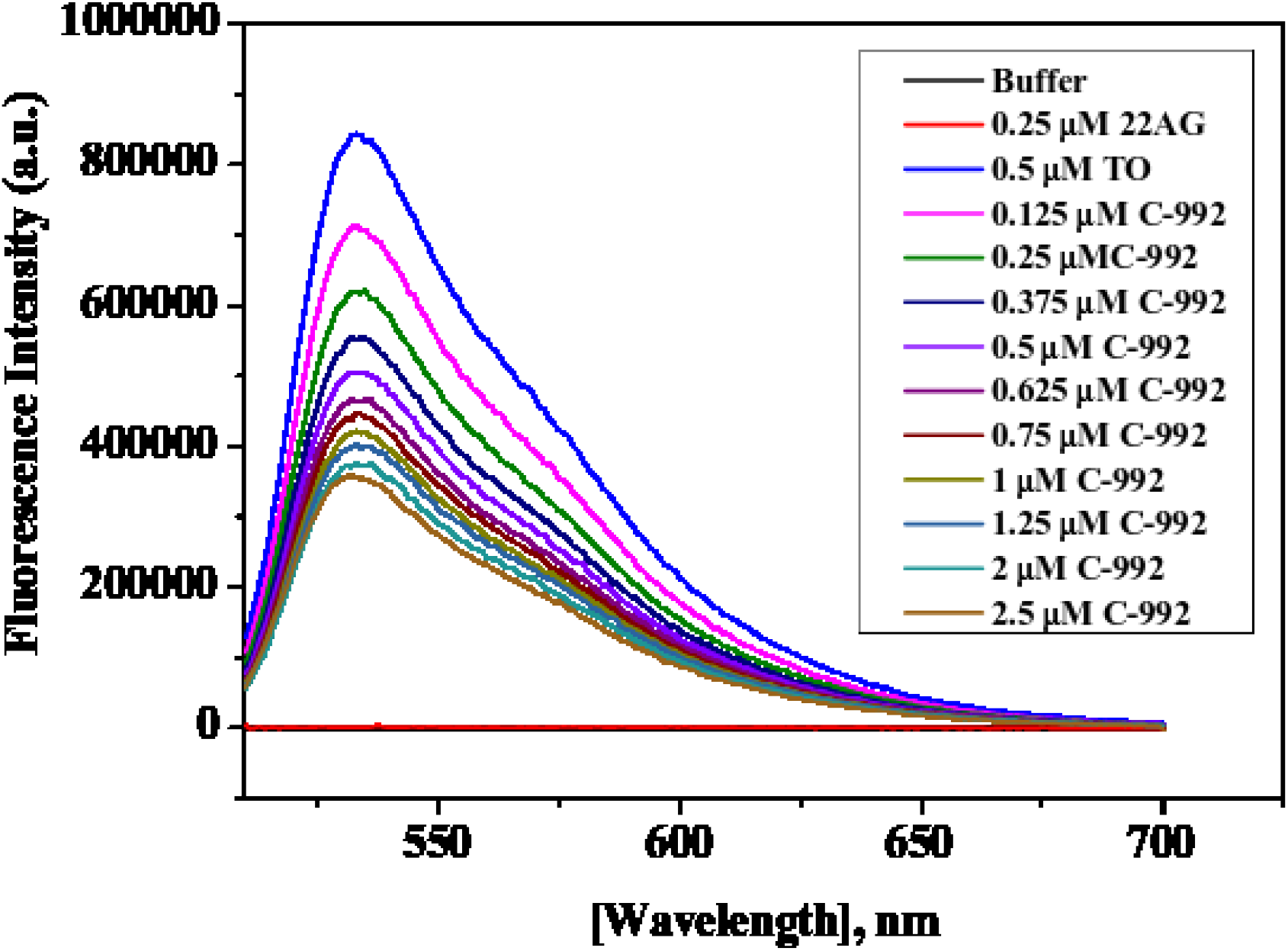
TO displacement by C-992 from TO-saturated 22AG G-quadruplex DNA

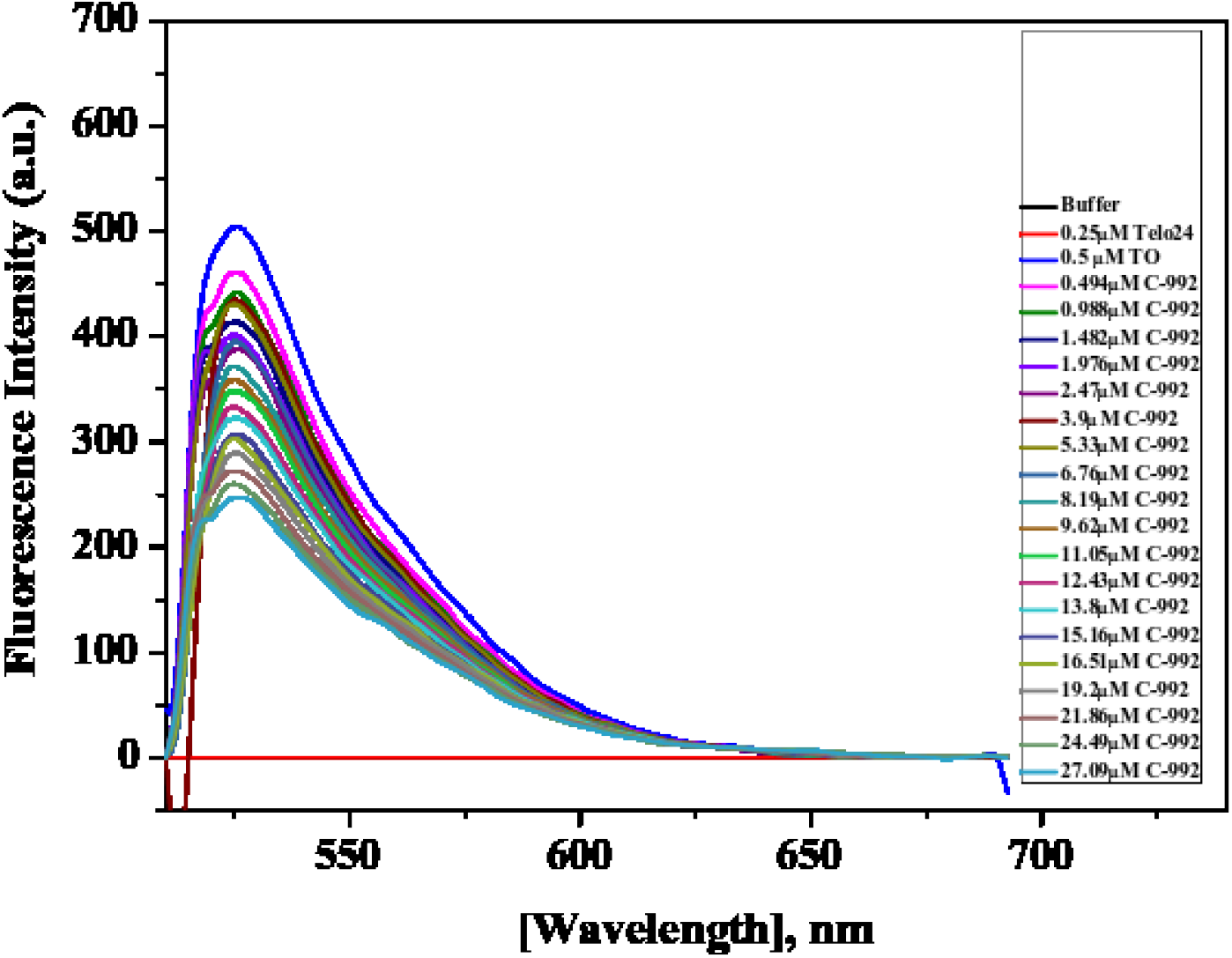
TO displacement by C-992 from TO-saturated Telo24 G-quadruplex DNA.

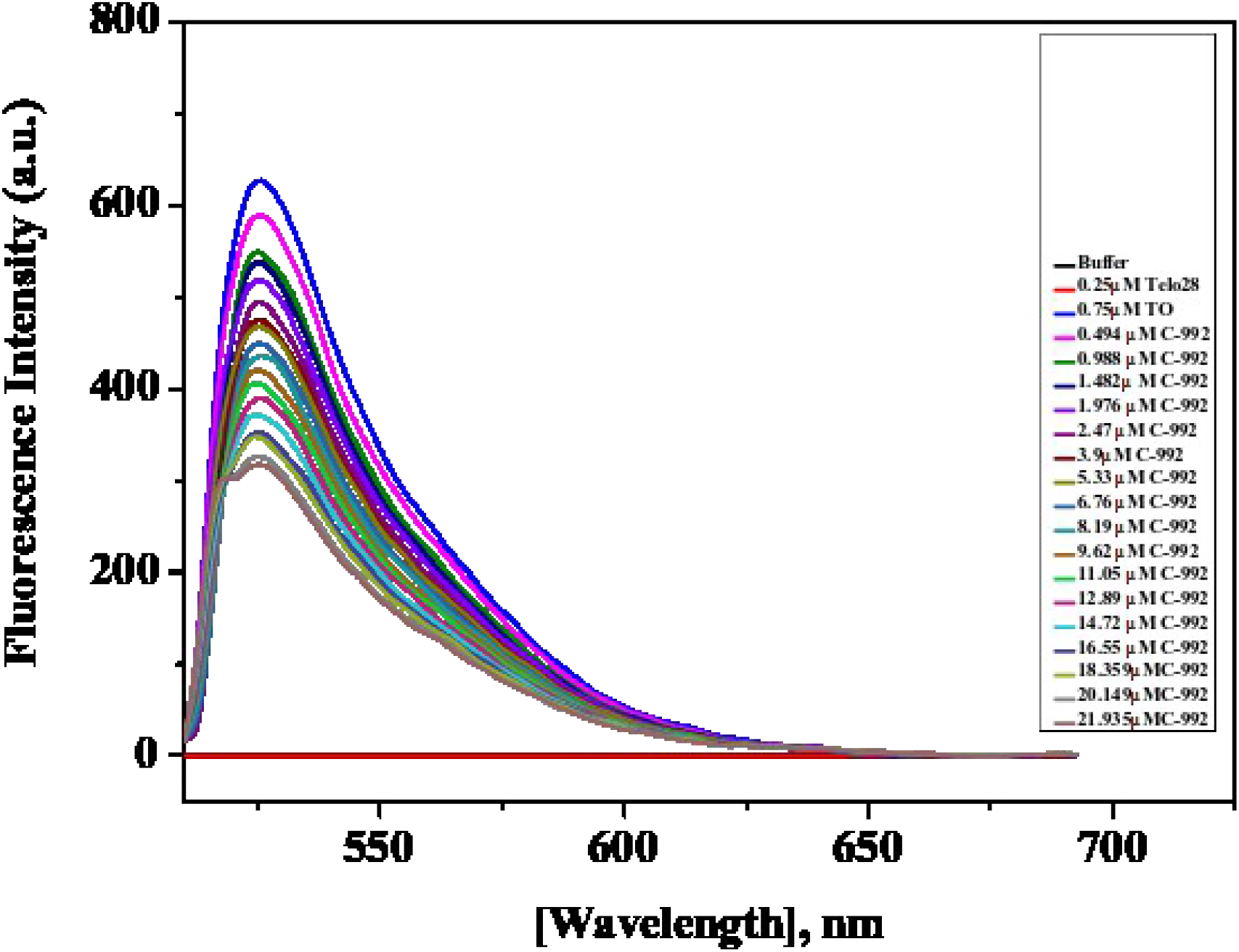
TO displacement by C-992 from TO-saturated Telo28 G-quadruplex DNA

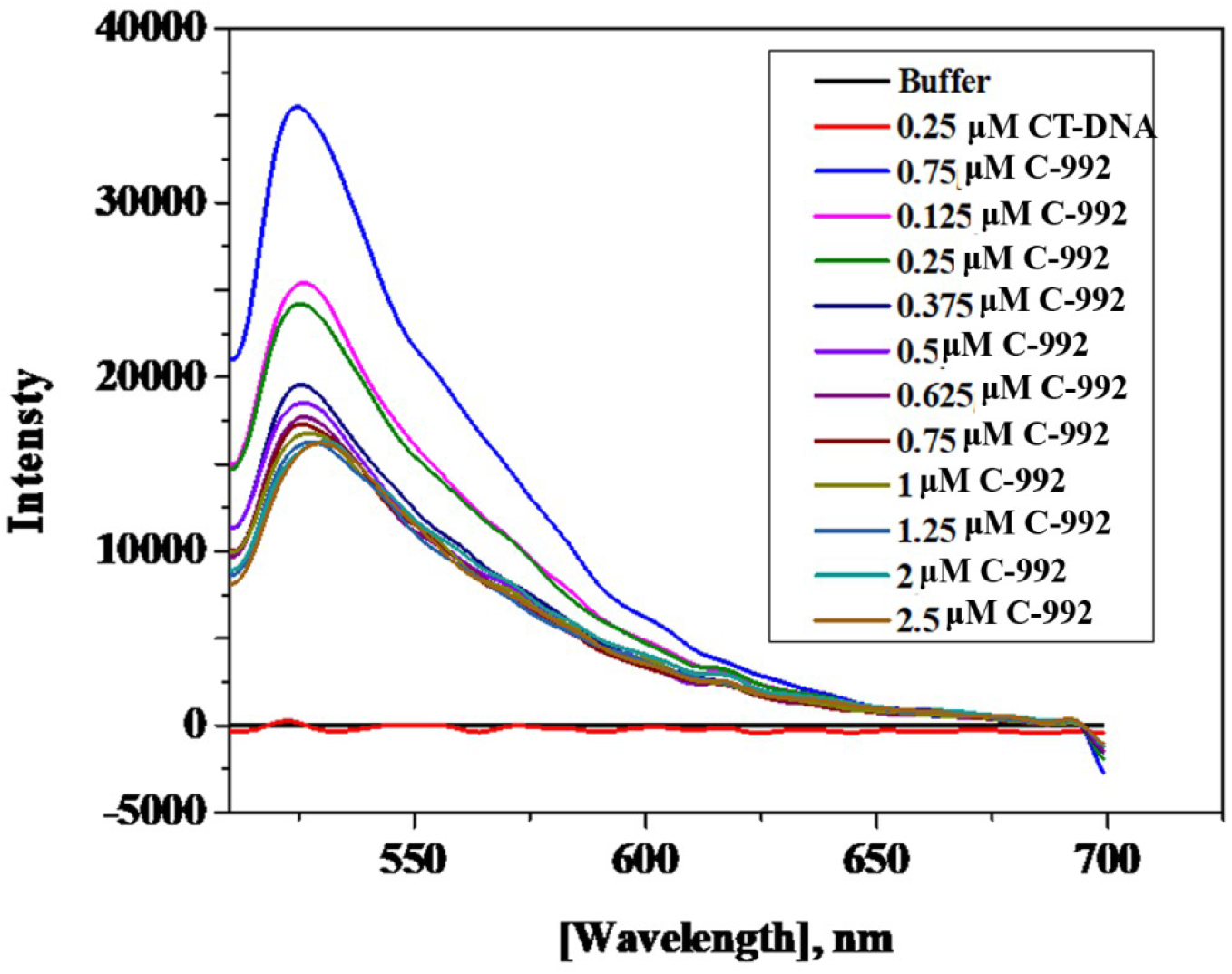
TO displacement by C-992 from TO-saturated CT-DNA

## Notes

### Competing Interest Statement

The authors have declared no competing interest.

